# Contiguous and accurate *de novo* assembly of metazoan genomes with modest long read coverage

**DOI:** 10.1101/029306

**Authors:** Mahul Chakraborty, James G. Baldwin-Brown, Anthony D. Long, J.J. Emerson

## Abstract

Genome assemblies that are accurate, complete, and contiguous are essential for identifying important structural and functional elements of genomes and for identifying genetic variation. Nevertheless, most recent genome assemblies remain incomplete and fragmented. While long molecule sequencing promises to deliver more complete genome assemblies with fewer gaps, concerns about error rates, low yields, stringent DNA requirements, and uncertainty about best practices may discourage many investigators from adopting this technology. Here, in conjunction with the platinum standard *Drosophila melanogaster* reference genome, we analyze recently published long molecule sequencing data to identify what governs completeness and contiguity of genome assemblies. We also present a hybrid meta-assembly approach that achieves remarkable assembly contiguity for both Drosophila and human assemblies with only modest long molecule sequencing coverage. Our results motivate a set of preliminary best practices for obtaining accurate and contiguous assemblies, a “missing manual” that guides key decisions in building high quality *de novo* genome assemblies, from DNA isolation to polishing the assembly.

## Introduction

*De novo* genome assembly is the process of stitching DNA fragments together into contiguous segments (contigs) representing an organism’s chromosomes (1). Until recently, genomes were often assembled using fragments shorter than 1,000 bp. However, such assemblies tend to be highly fragmented when they are generated using sequencing reads shorter than common repeats (1-4). Paired end short reads from different sized longer inserts can improve contiguity, but uncertainty of fragment length and the lack of sequencing between the insert ends makes resolving many repetitive structures challenging (5). Longer reads can circumvent this problem, even when such reads exhibit errors rates as high as 20% (5-9). Importantly, error-prone reads can be corrected, provided there is sufficient coverage and the errors are approximately uniformly distributed. Single molecule sequencing, like that offered by Pacific Biosciences (PacBio), meets these criteria with reads that are routinely tens of kilobases in length (5,10–12). While PacBio sequences have high error rates (~15%), errors are nearly uniformly distributed across sequences (5). With sufficient coverage, these sequences can be used to correct themselves (13). Assemblies using such correction are referred to as PacBio only assembly (14). Alternatively, hybrid assembly can be performed using a combination of noisy PacBio long molecules and high quality short reads (e.g. Illumina) (12,15).

Recently, the value of long molecule sequencing has been definitively demonstrated with the release of several high quality reference-grade genomes assembled from PacBio sequencing data (11,14). Indeed, the drosophila PacBio assembly closed gaps in the reference genome assembly (14), which is often considered the most contiguous metazoan genome assembly. Despite these successes, shepherding a genome project through the process of DNA isolation, sequencing, and assembly is still a challenge, especially for research groups for whom genomes are a means to another goal rather than the goal itself. For example, because high quality genome assembly relies upon long sequencing reads to bridge repetitive genomic regions(6,8,16,17) and high coverage to circumvent read errors (4,7,13), the stringent DNA isolation requirements (size, quantity, and purity) for PacBio sequencing (11) intended for genome assembly are higher than those typically employed. Moreover, at present, the low average read quality produced by PacBio sequencing causes coverage requirements to be at least 50-fold (5,14,18). This requirement, combined with the comparatively expensive sequencing, makes striking the right balance between price and assembly quality important. Exacerbating the problem is the fact that rediscovering the optimal approach for a genome project is itself expensive and time consuming. As a consequence of these challenges and uncertainties, many groups may opt out of a long molecule approach, or worse, sink scarce resources into an approach ill-suited for their goals because the consequences of many decisions involved in long molecule sequencing projects have not been synthesized.

In order to optimize a strategy for genome assembly we investigated the consequences of sample preparation (i.e. DNA isolation, quality control, shearing, library loading, etc.), assembly strategies, and properties of the data (i.e. read quality, length, and read filtering). We first evaluate strategies for assembling PacBio reads, and how they perform with differing amounts of sequence coverage. Then, we assess the contribution of read length and read quality to assembly contiguity. We also introduce quickmerge, a simple, fast, and general meta-assembler that merges assemblies to generate a more contiguous assembly. Additionally, we describe the protocols, quality-control practices, and size selection strategies that consistently yield high quality DNA reads required for reference grade genome assemblies. Our strategy is flexible enough to yield high quality assemblies using as little as 25X long molecule coverage or as much as >100X.

## Materials and methods

### Preparing high quality DNA library for long reads

#### Obtaining high quality, high molecular weight (HMW) genomic DNA

We used Qiagen’s Blood and Cell culture DNA Midi Kit for DNA extraction. As single molecule technologies (PacBio and Oxford Nanopore) do not require any sequence amplification step, a large amount of tissue is required to ensure enough DNA for library preparations that opt for no amplification (as is standard for genome assembly sequencing). For flies, 200 females or 250 males flies is sufficient for optimal yield (4060Mg DNA) from a single anion-exchange column. For other organisms, number of individuals need to be adjusted based on the tissue mass. A good rule of thumb is to keep the total amount of input tissue 100-150mg for optimal yield from each column.

To extract genomic DNA, 0-2 days old flies were starved for two hours, flash frozen in liquid nitrogen, and then ground into fine powder using a mortar and pestle pre-chilled with liquid nitrogen. The tissue powder was directly transferred into 9.5 ml of buffer G2 premixed with 38μl of RNaseA (100mg/ml) and then 250 μl (0.75AU) of protease (Qiagen) was added to the tissue homogenate. The volume of protease can be increased to 500 μl (1.5AU) to reduce the time of proteolysis. The tissue powder was mixed with the buffer by inverting the tube several times, ensuring that there were no large tissue clumps present in the solution. The homogenate was then incubated at 50°C overnight with gentle shaking (with 500μl protease, this incubation time can be reduced to 2 hours or less).

The next day, the sample was taken out of the incubator shaker and centrifuged at 5000xg for 10 minutes at 4°C to precipitate the tissue debris. The supernatant was decanted into a fresh 15ml tube. The little remaining particulate debris in the tube was removed with a 1 μl pipette. The sample was then vortexed for 5 seconds to increase the flow rate of the sample inside the column and then poured into the anion-exchange column. The column was washed and the DNA was eluted following the manufacturer’s protocol. Genomic DNA was precipitated with 0.7 volumes of isopropanol and resuspended in Tris buffer (pH 8.0). For storage of one week or less, we kept the DNA at 4°C to minimize freeze-thaw cycles; for longer storage, we kept the DNA at -20°C.

#### Shearing the DNA

1.5” blunt end needles (Jensen Global, Santa Barbara, CA) were used to shear the DNA. The needle size can be varied to obtain DNA of different length distribution: 24 gauge needles produces a size range of 24-50 kb. To obtain larger fragments, <24 gauge needles need to be used. For the DNA we have sequenced, up to 200ug of high molecular weight raw genomic DNA was sheared using the 24 gauge needle (Fig. 1). Additionally, we have also sheared DNA with 21, 22, and 23 gauge needles to demonstrate the size distribution they generate (supplementary Fig. 1). In brief, the entire DNA solution is drawn into a 1ml Luer syringe and dispensed quickly through the needle. This step is repeated 20 times to obtain the desired distribution of fragment sizes.

**Figure 1:**
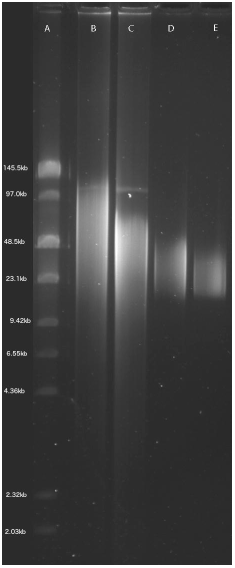
An example of correctly extracted and sheared DNA visualized using field inversion gel electrophoresis. The ladder is the NEB low range PFG marker (no longer produced). The lanes of the gel are as follows: (A) ladder, (B) unsheared DNA, (C) DNA sheared with a 24 gauge needle, (D) sheared DNA size selected with 15-50kb cutoff, (E) SMRTbell template library after 15-50kb size selection. From the gel, it is evident that there is a minimal ‘tail’ of DNA below ~15kb, the preferred size selection minimum.

#### Quality Control using FIGE

We verified the size distribution of unsheared and sheared genomic DNA using field inversion gel electrophoresis (FIGE), which allows separation of high molecular weight DNA. The DNA is run on a 1% agarose gel (0.5x TBE) with a pulse field gel ladder (New England Biolabs, Ipswich, MA). The gel is run at 4°C overnight in 0.5 x TBE. To avoid temperature or pH gradient buildup, a pump is used to circulate the buffer. The FIGE was run using a BioRad Pulsewave 760 and a standard power supply with the following run conditions:

> Initial time A: 0.6s, Final time B: 2.5 s, Ratio: 3, Run time: 8 h, MODE: 10, Initial time A: 2.5s, Final time B: 8s, Ratio: 3, Run time: 8 h, MODE: 11, Voltage: 135 V.

#### Library preparation

The needle sheared DNA is quantified with Qubit fluorometer (Life Technologies, Grand Island, NY) and NanoDrop (Thermo Scientific,,Wilmington, DE). Following quantification, 20 μg of sheared DNA was optionally run in four lanes of the Blue Pippin size selection instrument (Sage Science, Beverly, MA) using 15-50 kb as the cut-offs for size selection (Fig. 1). This optional size selection step increases final library yield at the cost of requiring more input DNA. This size selected DNA is then used to prepare a SMRTbell template library following PacBio’s protocol. A second round of size selection is performed on the SMRTbell template using a 15-50 kb cutoff to remove the smaller fragments generated during the SMRTbell library preparation step (Fig. 1). The second step minimizes the number of DNA fragments less than 15kb subjected to sequencing.

### DNA Sequencing

PacBio sequencing was conducted to demonstrate length distributions (*D. simulans* Fig. 2a) and evaluate the impact of library preparation on quality (Fig. 3), and was performed at the UCI High Throughput Core Facility using DNA isolated using the protocol described above. We note that the *D. simulans* reads were not used for assemblies reported here – all of our assemblies are constructed with publicly available *D. melanogaster* and *Homo sapiens* data (see Materials and methods). We sequenced one SMRTcell of *Drosophila* genomic DNA with the following conditions to obtain sequences with standard quality and length distribution: 10:1 polymerase to template ratio, 250 pM template concentration, and P6C4 chemistry. The movie time and other conditions were standard for RSII P6C4 chemistry. To demonstrate the tradeoff between yield and quality, we sequenced one SMRTcell each for polymerase:template ratios of 40:1,80:1,100:1 with template concentration held constant at 200pM, and one SMRTcell each with 300pM and 400pM template concentration with the polymerase:template ratio being held constant at 10:1.

**Figure 2.**
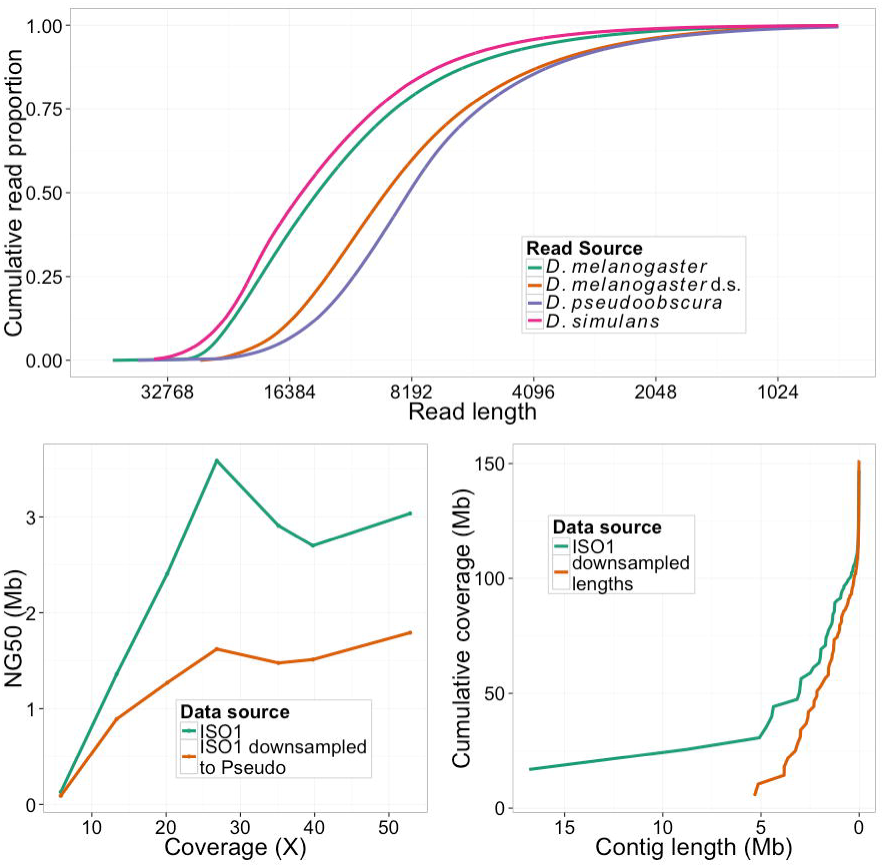
(a) The cumulative read length of various data sets, where *D. melanogaster* refers to the original ISO1 data set, *D. pseudoobscura* refers to a publicly available *D. pseudoobscura* dataset with a shorter average read length, *D. melanogaster* d.s. refers to the *D. melanogaster* data, downsampled to have read lengths resembling the *D. pseudoobscura* dataset, and *D. simulans* is a *D. simulans* dataset sequenced using our DNA preparation technique. (b) A plot of NG50 versus coverage of hybrid assemblies, as in Fig. 5. This plot depicts the effect of reduced read length on NG50, while holding read quality and coverage constant. (c) Cumulative contig length distribution of 53X of PacBio only assemblies created with the original ISO1 reads and the ISO1 reads downsampled to resemble *Pseudoobscura.* Contig lengths in the shorter/downsampled reads assembly are considerably shorter than the contigs in the original reads assembly.

**Figure 3.**
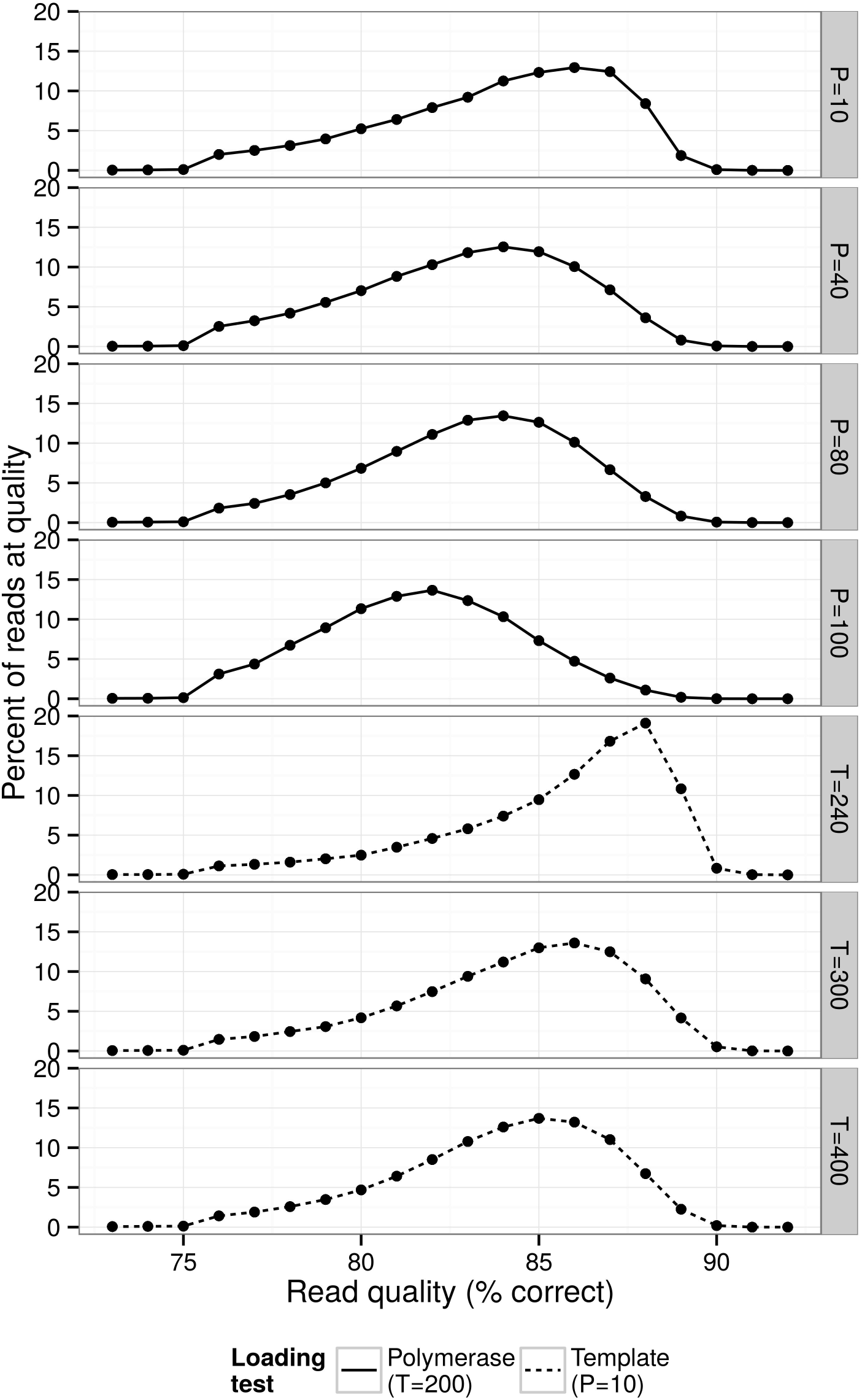
The distribution of read quality in sequencing runs performed at the UCI genomics core using our DNA preparation technique. “P” here refers to polymerase loading during sequencing (the proportion of polymerase to template, where 10 would indicate a 10:1 ratio of polymerase to template), while “T” refers to template loading concentration during sequencing (in picomolarity).

### PacBio only Assembly

For PacBio sequences, the assembly pipeline is divided into three parts: correction, assembly, and polishing. Correction reduces the error rate in the reads to 0.5-1% (14), and is necessary because reads with a high (~15%) error rate are extremely difficult to assemble (17). Correction is facilitated by high PacBio coverage, which allows the error corrector to successfully ‘vote out’ errors in the PacBio reads. For self correction, we used the PBcR pipeline (14) as implemented in wgs8.3rc1 which, by default, corrects the longest 40X reads. The second step involves assembling the corrected reads into contigs. We used the Celera assembler (17), included in the same *wgs* package, for assembly. A third optional step involves polishing the contigs using *Quiver* and *Pilon* (19, 20), which brings the error rate down to 0.01% or lower. All of the assemblies described in this paper were generated with the same PBcR command and spec file (commands and settings, Supplementary materials).

For PacBio only assembly of *D. melanogaster* ISO1 sequences, we used a publicly available PacBio sequence dataset which was generated using the standard P5C3 chemistry. A complete description of this data is available in Kim et al. (11). We chose the *D. melanogaster* dataset for our experiments and simulations because *D. melanogaster* is widely used in genetics and genomics research and its reference sequence (release 5.57,http://www.fruitfly.org) is one of the best, if not the best, eukaryotic multicellular genome assemblies in terms of assembly contiguity. This is true for both the PacBio generated assembly (21Mb contig N50)^13^ and the Sanger assembly (23Mb scaffold N50) of ISO1. The remarkable contiguity of these assemblies becomes more tangible when the theoretical limits of *D. melanogaster* chromosome arms’ lengths are considered (21): N50 of both assemblies lie very close to the theoretical maximum N50 (~28Mb). This high quality assembly serves as a reference for evaluating assemblies presented here.

We evaluated assembly qualities using the standard assembly statistics (average contig size, number of contigs, assembled genome size, N50, etc.) using the *Quast* and *GAGE* (22, 23) packages.

### Hybrid Assembly

PacBio only assembly of high error, long molecule sequences depends upon redundancy between the various low quality reads to ‘vote out’ errors and identify the true sequence in the sequenced individual. An alternative approach to this problem is to use known high quality sequencing reads to correctly call the bases in the sequence, and then to use PacBio reads to identify the connectivity of the genome. In order to achieve the best possible assembly results, we tested several different hybrid assembly pipelines before choosing *DBG2OLC* and *Platanus* (24). In our early tests, the next highest performing hybrid assembler, a combination of *ECTools* (25) and *Celera,* achieved a highest N50 of 616kb in *Arabidopsis thaliana* using 19 SMRT cells of data (25); in contrast, using 20 SMRT cells of the same data, the *DBG2OLC* and *Platanus* pipeline produced an N50 of 4.8Mb. We aslo tested the alternative error corrector, *LorDEC* (26), along with the *Celera* assembler, but found that the Lordec-corrected Celera assembly of our standard *D. melanogaster* dataset (26X of PacBio data and 67.4X of Illumina data) produced an NG50 of only 109KB. Consequently we adopted *DBG2OLC* as our choice for hybrid assembly. The hybrid error corrector *LSC* (27) was not tested in detail due to the fundamental similarity of its assembly approach to that of *ectools* and *Lordec,* though preliminary data (not shown) did not indicate superior read correction by *LSC* versus these other approaches. Using the standard 67.4X of Illumina data discussed above and 26X of PacBio data, we compared *DBG2OLC* runs using three different De Bruijn graph assemblers: *SOAP* (28), *ABySS* (29), and *Platanus.* The NG50s for the three assemblies were, respectively, 2.43Mb, 0.167Mb, and 3.59Mb. Based on this result, we chose to use *Platanus* for the remainder of the assemblies.

We used the pipeline recommended by *DBG2OLC* (30) to perform hybrid assemblies. In this pipeline, we used *Platanus* to perform De Bruijn graph assembly on the Illumina reads. We used 8.36 Gb (67.4X) of Illumina sequence data of the ISO1 *D. melanogaster* inbred line generated by the DPGP project (31) to generate a De Bruijn graph assembly using *Platanus.* We used *DBG2OLC* to align our PacBio reads to the De Bruijn graph assembly to produce a ‘backbone’, then, according to the DBG2OLC standard pipeline, used the backbone to generate the consensus using the programs *BLASR* (32) and *PBDagCon* (https://github.com/PacificBiosciences/pbdagcon). As with the PacBio only assemblies above, we evaluated assembly quality using the *Quast* and *GAGE* packages.

### Assembly merging

Hybrid assembly and PacBio assembly were merged using a custom C** program (Fig. 4A, available at https://github.com/mahulchak/quickmerge). The program takes two fasta files (containing contigs from a PacBio only assembly and contigs from a hybrid assembly) as inputs and splices contigs from the two assemblies together to produce an assembly with higher contiguity. As the two assemblies used for merging come from the same genome, gaps in one assembly can be bridged using homologous sequences from the other assembly. The first stage of the assembly merging process involves correctly aligning the homologous sequences (contigs), which in the second stage are exchanged at the sequence gaps so that the part of the sequence with the gap is replaced with a contiguous sequence from the other assembly. The program MUMmer (33) is used to find the correct alignment between the assemblies and assembly merging is handled by *quickmerge*.

**Figure 4:**
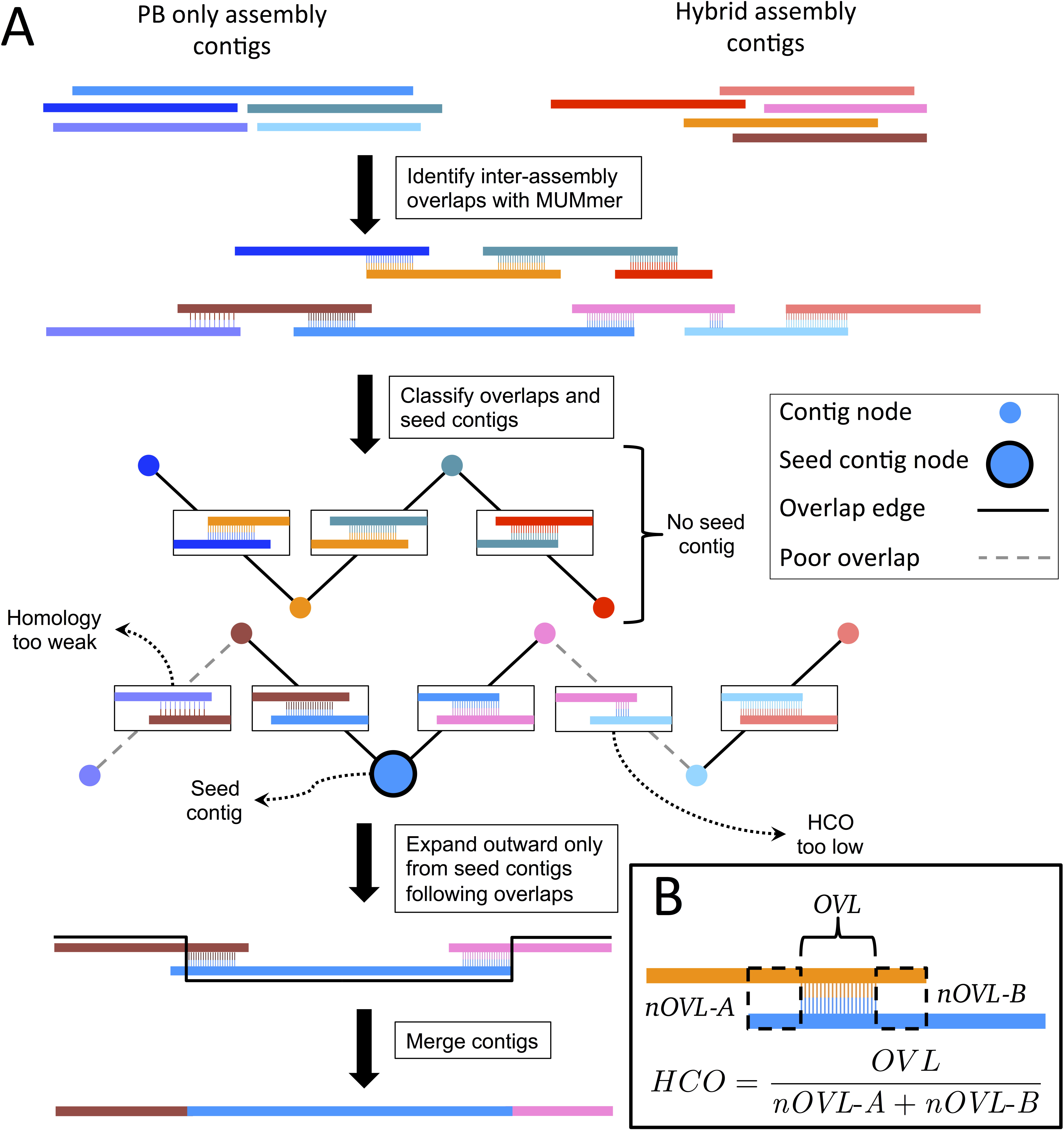
A) A diagram representing the algorithm employed by quickmerge to improve genome contiguity. (A) MUMmer is used to identify overlaps between the two assemblies. High confidence overlaps (HCOs) identified by MUMmer will be the primary signal to quickmerge that two contigs should be joined. Quickmerge clusters contigs according to HCOs. Quickmerge identifies seed contigs (contigs in a cluster above a certain size and HCO), and identifies a path that connects it to all other contigs in its cluster by walking from one contig to the next, only stepping to the next contig if the quality of the HCO between the current and next contigs is above the set thresholds. Once the graph connecting available contigs to the seed contig has been constructed, the contigs in the graph are spliced together, with the “Donor” genome’s content preferred over the “acceptor” genome. B) Description of the HCO parameter. HCO represents the ratio between overlapping aligned and overlapping unaligned parts between two contigs.

First, the program *MUMmer* (33) is used to compute the unique alignments between the contigs from the two assemblies, one of which is used as the reference, or donor, assembly and the other is used as the query, or acceptor, assembly. Distinction between the two assemblies is important because, as described below, the user may choose the more reliable (with fewer errors) of the two assemblies to bridge gaps in the other assembly. Accurate merging occurs when homology between two sequences is high; conversely, pairing between non-homologous regions leads to incorporation of incorrect sequences. Hence, identification of the true homologous pairing is necessary for error-free sequence merging. Presence of repeats may complicate the situation, but the problem can largely be overcome if the two aligned sequences containing repeats come from the same genome and only the unique best alignments are considered. To obtain the unique best alignment between the reference and the query assembly, spurious matches introduced by gene duplications and repeats are removed using the delta-filter utility (with –r and –q options) of the MUMmer package.

Following the repeat filtering step, the alignments are partitioned using a scoring metric called high confidence overlaps (HCO) (Fig. 4B). The program identifies HCOs by dividing the total alignment length between contigs by the length of unaligned but overlapping regions of the alignment partners (Fig. 4B). The metric was chosen under the assumption that the length of the overlapping but unaligned portion between the two sequences relative to the length of the overlapping and aligned parts is high for two non-homologous sequences. After the alignment partitioning is done based on a HCO cutoff, only the contig alignments above the HCO cutoff are kept for assembly merging.

For fly assemblies, we found that an HCO value of 1.5 to an appropriate default for assembly merges. This cutoff can be increased further, as we did for merging human assemblies. The tradeoff is that increasing HCO cutoff will gradually deplete the pool of “homologous” alignments, thereby leading to a reduction in merging events. Thus, the “HCO” parameter controls merging sensitivity at the cost of increased false positives: the higher the HCO parameter value, the more stringent is the cutoff for HCO selection.

The next step involves searching and ordering the contigs that will be merged. To accomplish that, by default *quickmerge* assigns nodes in the HCO alignment graph with even higher HCO values (>5.0) and reference sequences exceeding a length cutoff (1Mb) as anchor nodes. The high HCO and the length cutoff are used here to ensure that subsequent searches for contigs for merged contig extension do not begin at spurious alignment nodes. Following the assignment of the anchor nodes, a greedy search is initiated on both the left and the right sides (5’ and 3’ of the reference contig) of the anchor node, in order to find the longest unbroken path through the HCO nodes.

In other words, *quickmerge* looks for contigs that connect two adjacent HCO nodes in the graph and this process is continued until no contig can be found to connect two HCO nodes (e.g. a genomic region where both assemblies are broken). For the search, each contig is used only once to connect two HCO nodes, so once a contig from the HCO alignment pool has been used, it is removed from the alignment pool. Query contigs that are completely contained within a reference contig are also removed from the final merged assembly to prevent sequence duplication in the merged assembly.

In the final step, the ordered chain of contigs found in the previous step is joined by swapping portions of the reference assembly into the query assembly in a manner that maximizes retention of sequences from the reference assembly (Fig. 4A). Gap filling within the query assembly occurs as a byproduct of this replacement of sequences; in this way, the process resembles genome editing using homologous recombination.

For coverages of 40X, 53X, 62X, and 77X, merged assemblies were generated using the PacBio only assembly and their corresponding hybrid assemblies. For the 99x and 121x (all reads) SMRTcells datasets, the PacBio only assemblies were merged with the hybrid assembly obtained from the 77X SMRTcells dataset. All hybrid assemblies used for merging were generated without downsampling by read length or quality. The time to merge was limited only by the time required to run *MUMmer,* as *quickmerge* runs in less than 30 seconds on *Drosophila*-sized genomes, and requires less than 2GB of memory.

### Downsampling

We used a number of different downsampling schemes on the *D. melanogaster* data: first, we randomly downsampled the data by drawing a random set of SMRTcells of data from the entire set of 42 SMRTcells; second, from those datasets, we downsampled the longest 50% and 75% of the reads. next, we downsampled the *D. melanogaster* data to match the read length distributions of PacBio reads from a pilot *Drosophila pseudoobscura* genome project that was produced using a standard protocol without aggressive size selection (generously made available by Stephen Richards). Finally, we downsampled based on read quality to test the effect of read quality on assembly contiguity. Please see the supplementary text for more details.

## Results

### DNA isolation for long reads

As the remainder of the paper will show, read length is an important determinant of genome assembly contiguity. We identified simple and consistent method for isolation of large genomic DNA fragments necessary for PacBio sequencing to achieve long reads. The existing alternative method used for DNA isolation to generate the published PacBio Drosophila assembly involved DNA extraction by CsCl density gradient centrifugation and g-Tube (Covaris, Woburn, MA) based DNA shearing (11). CsCl gradient centrifugation is a time-consuming method that requires expensive equipment that is not routinely found in most labs. Additionally, g-Tubes are expensive, require specific centrifuges, and are extremely sensitive to both the total mass of DNA input and to its length. We circumvented these problems by using a widely available DNA gravity flow anion exchange column extraction kit in concert with a blunt needle shearing method (32). Because the DNA fragment size distribution is so important, field inversion gel electrophoresis (FIGE) is an essential quality control step to validate the length distribution of the input DNA (Fig. 1) (see Methods for details). Sequences generated from libraries constructed from this isolation method are comparable to or longer than the published Drosophila PacBio reads (11) (Fig. 2a). The length distribution of the input DNA can potentially be improved further by using wider gauge needles that generate even longer DNA fragments (supplementary Fig. 1).

### Long read assembly

PacBio self correction has been used to assemble the *D. melanogaster* reference strain (ISO1) genome so contiguously that most chromosome arms were represented by fewer than 10 contigs(14). This assembly was generated by using the PBcR pipeline (14) and 121X (15.8 Gb), or 42 SMRTcells’ worth, of PacBio long molecule sequences(11). However, currently, such high coverage may be too expensive for many projects, especially when the genome of the target organism is large. Consequently, we set out to determine how much sequence data is required to obtain assemblies of desired contiguity. We first selected reads from 15, 20, 25, 30, and 35 randomly chosen SMRTcells (40X, 53X, 62X, 77X, and 99X assuming a genome size of 130×10^6^ bp – coverages calculated by dividing total bases of sequence data by total bases in genome) from the 42 SMRTcells of ISO1 PacBio reads (11). Our sampling method was inclusive and additive: for example, to obtain 20 SMRTcells, we took the 15 previously randomly chosen SMRTcells and then added 5 more randomly selected SMRTcells to it. We then assembled these datasets using the PBcR pipeline. As shown in Fig. 5, the contig NG50 (NG50; G =130×10^6^ bp) continues to improve across the entire range of coverage. At extremely high coverage (121x), the NG50 surges again, approaching the theoretical N50 limit of *D. melanogaster* genome (21). Notably, despite the extreme contiguity of these sequences, we are still discussing complete contigs, not scaffolds with gaps.

**Figure 5.**
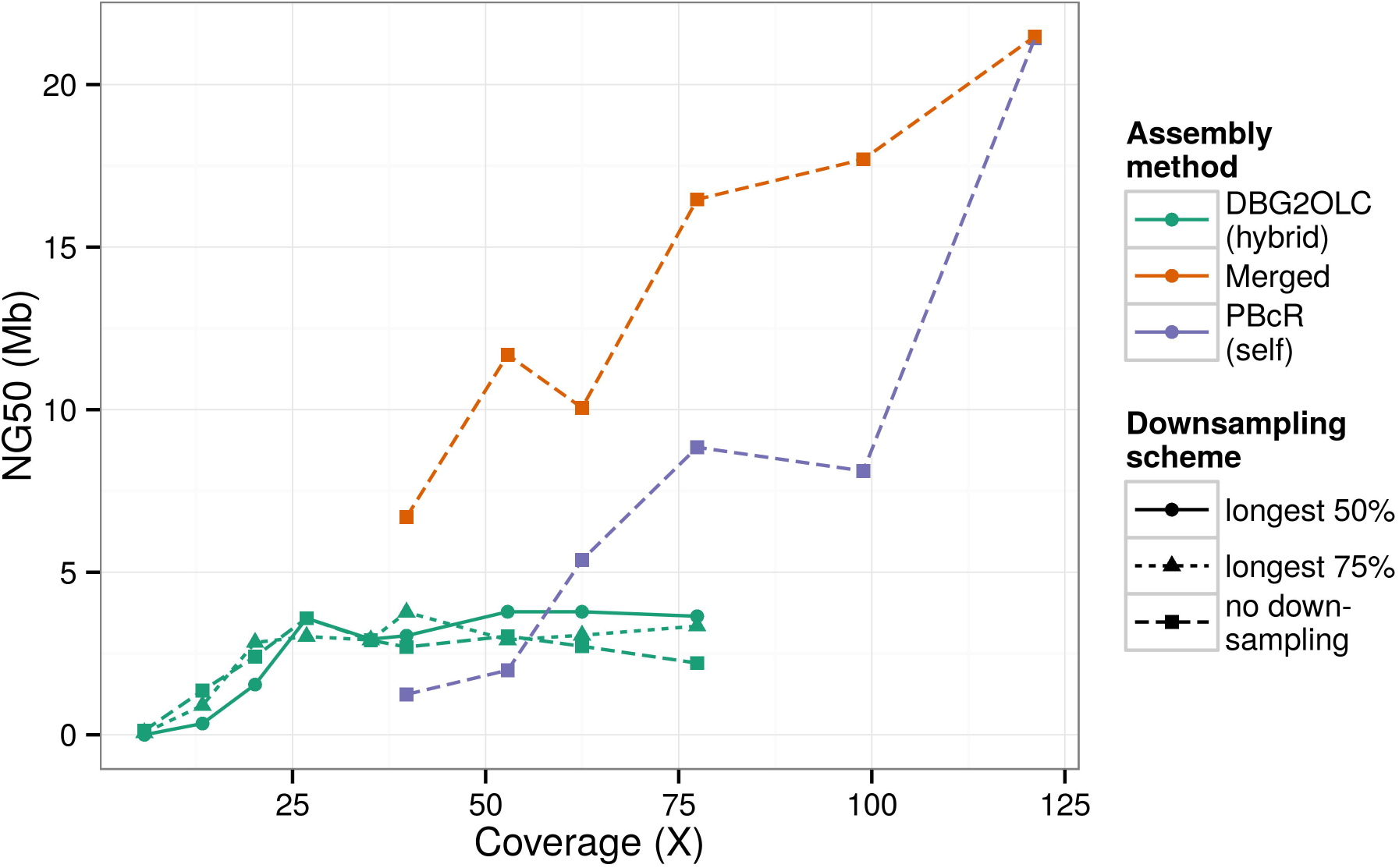
The NG50 of *D. melanogaster* assemblies produced using a variety of data sets. NG50 here is the contig size such that at least half of the 130Mb *D. melanogaster* genome (65Mb) is contained in contigs of that size or larger. “longest 50%” and “longest 75%”, respectively, refer to datasets in which only the longest 50% or 75% of the available reads have been used. The coverage listed on the x-axis in this case refers to the total amount of available data (before downsampling by length).

### Hybrid assembly

As Fig. 5 makes clear, PacBio only assembly leads to relatively fragmented genomes at lower coverage (Fig. 5), we investigated whether another assembly strategy could perform better with similar amounts of long molecule data. We chose DBG2OLC (29) for its speed and its ability to assemble using less than 30X of long molecule coverage (*cf.* PacBio only methods, which typically require higher coverage (5)). DBG2OLC is a hybrid method, which uses both long read data and contigs obtained from a De Bruijn graph assembly. We used contigs from a single Illumina assembly generated using 67.4X of Illumina paired end reads (30). As shown in Fig. 5, the assembly NG50 increases dramatically as PacBio coverage increased, plateauing near 26X. Beyond this point, NG50 remained relatively constant. Alignment of the test assemblies to the ISO1 reference genome showed that some of the contiguity in the 26X hybrid assembly without downsampling was due to chimeric contigs (ie contigs that possess non-syntenic misjoins), and that these errors are fixed as coverage increases (supplementary Fig. 2–3). Chimeras were also absent when only the longest 50% or 75% of reads from the 26X dataset were used.

To measure the impact of read length on hybrid assembly contiguity, we downsampled the datasets by discarding the shortest reads such that the resulting datasets contained 50% and 75% of initial total basepairs of data. We then ran the same assembly pipelines using these downsampled datasets and compared to the assemblies constructed from their counterparts that were not downsampled. Our downsampling shows that with high levels of PacBio coverage (> 50x), modest gains in assembly contiguity can be obtained by simply discarding the shortest reads (Fig. 5, green lines). Our hybrid assembly results indicate that improvements in contiguity above 30X are modest, though hybrid assemblies remain more contiguous than PacBio only assemblies up until above 60X coverage. For projects limited by the cost of long molecule sequencing, a hybrid approach using ~30X PacBio sequence coverage is an attractive target that minimizes sequencing in exchange for modest sacrifices in contiguity that are in any event available only at higher coverages.

### Assembly merging

With modest PacBio sequence coverage (≤50X), hybrid assemblies are less fragmented than their self corrected counterparts, but more fragmented than self corrected assemblies generated from higher read coverage (Fig. 5). Despite this, for lower coverage, many contigs exhibit complementary contiguity, as observed in alignments (e.g. Supplementary Fig. 4a) between a PacBio only assembly (53X reads; NG50 1.98 Mb) and a hybrid assembly (longest 30X from 53X reads; NG50 3.2 Mb; not featured in Fig. 5). For example, the longest contig (16.8 Mb) in the PacBio only assembly, which aligns to the chromosome 3R of the reference sequence (Supplementary Fig. 4c), is spanned by 5 contigs in the hybrid assembly (Supplementary Fig. 4b). This complementarity suggests that merging might improve the overall assembly.

We first attempted to merge the hybrid assembly and the PacBio only assembly using the existing meta assembler minimus2 (35), but the program often failed to run to completion when merging a hybrid assembly and a PacBio only assembly, and when it did finish, the run times were measured in days. We therefore developed a program, *quickmerge,* that merges assemblies using the MUMmer (33) alignment between the assemblies. Assembly contiguity improved dramatically when we merged the above hybrid and PacBio only assemblies (assembly NG50 9.1 Mb; supplementary Fig. 5); however, assembly contiguity can also be increased with false contig joining. To investigate whether merging leads to false joins or introduces assembly errors at the splice junctions, we investigated the result of merging at base pair resolution for the longest merged contig in the aforementioned assembly.

The longest contig (27.9 Mb) in the merged assembly, which aligns to chromosome arm 3R of the reference sequence (supplementary Fig. 6), was longer than the longest 3R contig in the PacBio assembly based on 42 SMRTcells (25.4Mb)(14) (supplementary Fig. 6). The increased length resulted from closing of gaps present in the published PacBio assembly (supplementary Fig. 6) (14). All joined contigs map to the chromosome arm 3R in the correct order; we take this as evidence that *quickmerge* does not incorporate spurious sequences or large scale misassemblies

Nonetheless, small scale misassemblies could still be introduced at the splice junctions. To check for such errors, we manually inspected a high resolution dot plot between the merged contig and the 3R reference sequence. A total of 18 regions were found where the merged contig differed from the reference sequence (supplementary Table 2). The affected regions ranged from 3bp to 20 kb and involved sequence insertion, deletion, and duplication. All identified misassemblies had a buried Pacbio coverage of 15 or higher, indicating that misassemblies were due not to lack of coverage, but some other factor (for example, repetitive regions of the genome). For buried coverage calculations, reads are mapped to the genome, and only mapped regions supported by 2kb unbroken read coverage on both sides are counted towards buried coverage, ensuring any feature exhibiting buried coverage is strongly supported by the reads overlapping it.

That said, such discordance between the merged contigs and the reference could have been carryover assembly errors from the hybrid and PacBio only assemblies that were used for merging. Indeed, 11 of the 18 errors in the merged contigs came from the PacBio only assembly, whereas the rest came from the hybrid assembly. Additionally, sequences 201bp in length from each of the 29 splice joints (break point is the101th base pair, see Supplementary text) from the aforementioned merged assembly were aligned to the reference sequence. None of the sequences revealed any misassemblies introduced by the merging process. Thus, for this dataset, the *quickmerge* approach splices and merges contigs accurately without introducing any new assembly errors.

This indicates that the contiguity of even high coverage PacBio only assemblies can be increased by the addition of inexpensive Illumina reads, and gaps in hybrid assembly can be closed by PacBio only assembly even when the PacBio only assembly quality is suboptimal.

### Assessment of assembly quality

We assessed assembly quality using the *Quast* software package (22) and the quality assessment scripts used in the GAGE study (23). We confined our assessment to assemblies related to application of the quickmerge meta assembler, leaving the assessment of PBcR and DBG2OLC assemblies to their respective publications (14,29). Quast quantifies assembly contiguity and additionally identifies misassemblies, indels, gaps, and substitutions in an assembly when compared to a known reference. We found that, compared to the *D. melanogaster* reference, all assemblies had relatively few errors, with the primary difference among the assemblies being genome contiguity (NG50). Hybrid assemblies tended to have fewer assembly errors than PacBio only assemblies: the total number of misassemblies and the total number of contigs with misassemblies tended to be higher in PacBio only assemblies compared to hybrid assemblies. Still, PacBio only assemblies tended to have slightly fewer mismatched bases compared to the reference, and slightly fewer small indels. Merged assemblies, being a mix of PacBio only and hybrid assemblies, tended to have intermediate *Quast* statistics; however, the merged assemblies improved upon the source assemblies in terms of misassemblies and misassembled contigs. Overall, the rate of mismatches was low at an average (across all assemblies) of 47 errors per 100kb (Supplementary Table 1, Supplementary Fig. 8). Mismatches and indels can be further reduced using existing programs, such as *Quiver* (19). We used *Quiver* to polish all non-downsampled hybrid, self, and merged assemblies that used at least 40X of data. After *Quiver,* the average mismatch rate of the selected assemblies decreased from 24 per 100kb to 15, while the average indel rate decreased from 180 per 100kb to 32 (Supplementary Fig. 9). We also performed post-Quiver polishing on these selected assemblies using Illumina data via the *Pilon* program (20). Pilon polishing further reduced the average indel rate per 100kb from 32 to 16 (Supplementary Fig 10).

One concern generated by the pre-polished assemblies was that their N50s were high, but their corrected N50s (23) after accounting for errors were low; however, Quiver and Pilon polishing dramatically improved the corrected N50s of the assemblies, indicating that the low corrected N50 values were due to small local errors that were easily resolved by polishing. The average corrected N50 before polishing was 67kb, while the average corrected N50 after polishing was 530kb. It is evident from the corrected N50s that the first polishing step, Quiver, was responsible for most of the change in corrected N50 (Supplementary Fig. 11).

### Size selection and assembly contiguity

Long reads generated by library preparation with aggressive size selection (11) can generate extremely contiguous and accurate *de novo* assemblies (14). Unfortunately, some DNA libraries with less stringent size selection produce considerably shorter reads (Fig. 2a). Longer reads are predicted to generate more contiguous genomes (6,7). We tested this hypothesis by assembling genomes using randomly sampled whole reads (see Materials and Methods) from the ISO1 dataset to simulate a read length distribution comparable to, but slightly longer than what is typical when size selection is not aggressive. Due to the long read length distribution of the ISO1 dataset relative to the shorter target distribution above, a maximum of 53X of ISO1 data could be sampled.

Consistent with the theoretical prediction that, all else being equal, shorter reads produce more fragmented assemblies(6,7), reads from the downsampled 53X ISO1 data produced a PacBio only assembly with an NG50 of 1.38 Mb, which is shorter than the NG50 (1.98 Mb) of the assembly from the same amount of ISO1 long read data (Fig. 2c). In addition, nearly all long contigs present in the original 53X assembly are fragmented in the assembly from the shorter reads (Supplementary Fig. 13), although the amount of sequence data (53X) used to build the assemblies is the same.

For hybrid assembly, the shorter dataset also produced significantly less contiguous assemblies, consistent with predictions from theory (7) (Fig. 2b). The NG50 achieved with 26X coverage of the shorter dataset was 1.62Mb, compared to an NG50 of 3.58Mb with the original ISO1 data. This is consistent with the PacBio only result – longer read lengths lead to higher assembly contiguity. Thus, a library preparation procedure that aggressively size selects DNA is crucial in delivering long contigs.

### The effects of read quality on assembly

As with reduction in read length, increased read errors are predicted to worsen assembly quality because noisier reads increase the required read length and coverage to attain a high quality assembly (9,13). When a PacBio sequencing experiment is pushed for high yield through either high polymerase or template concentration, the data exhibits lower quality scores (Fig. 3). Thus, with equal coverage and read length distribution, reads with higher error rates should result in a more fragmented assembly. To measure this effect, we partitioned the ISO1 PacBio read data into three groups with equal amounts of sequence without changing the read length distribution (see methods) (Supplementary Fig. 14). For the first two groups, the data was split in half, with one half comprising the reads from the bottom 50% of phred scores and the other comprising the top 50%. The third dataset was generated by randomly selecting 50% of the reads in the full dataset. We then performed PacBio-only and hybrid assemblies with these data.

Low read quality had a particularly dramatic effect on assembly by self correction (Fig. 6): the high quality and randomly sampled reads produced substantially better assemblies (6.23 Mb and 6.15 Mb, respectively) than the assembly made from low quality reads (NG50 146 kb). Hybrid assembly contiguity was far more robust to low quality reads (Fig. 6: NG50 of 3.1Mb for the high quality reads, 2.5Mb for the unfiltered reads, and 2.2Mb for the low quality reads), showing only moderate variation amongst different quality datasets. Throughout this study, we avoided altering the settings from their default states in the various assemblers used in order to do fair comparisons; however, in this case, we chose to also run PBcR in ‘sensitive’ mode to see if it would improve contiguity when data quality is low. We found assembly contiguity was improved (NG50=4Mb), but was still lower than the assembly generated from unselected reads without the sensitive parameters (NG50=6.23Mb).

**Figure 6.**
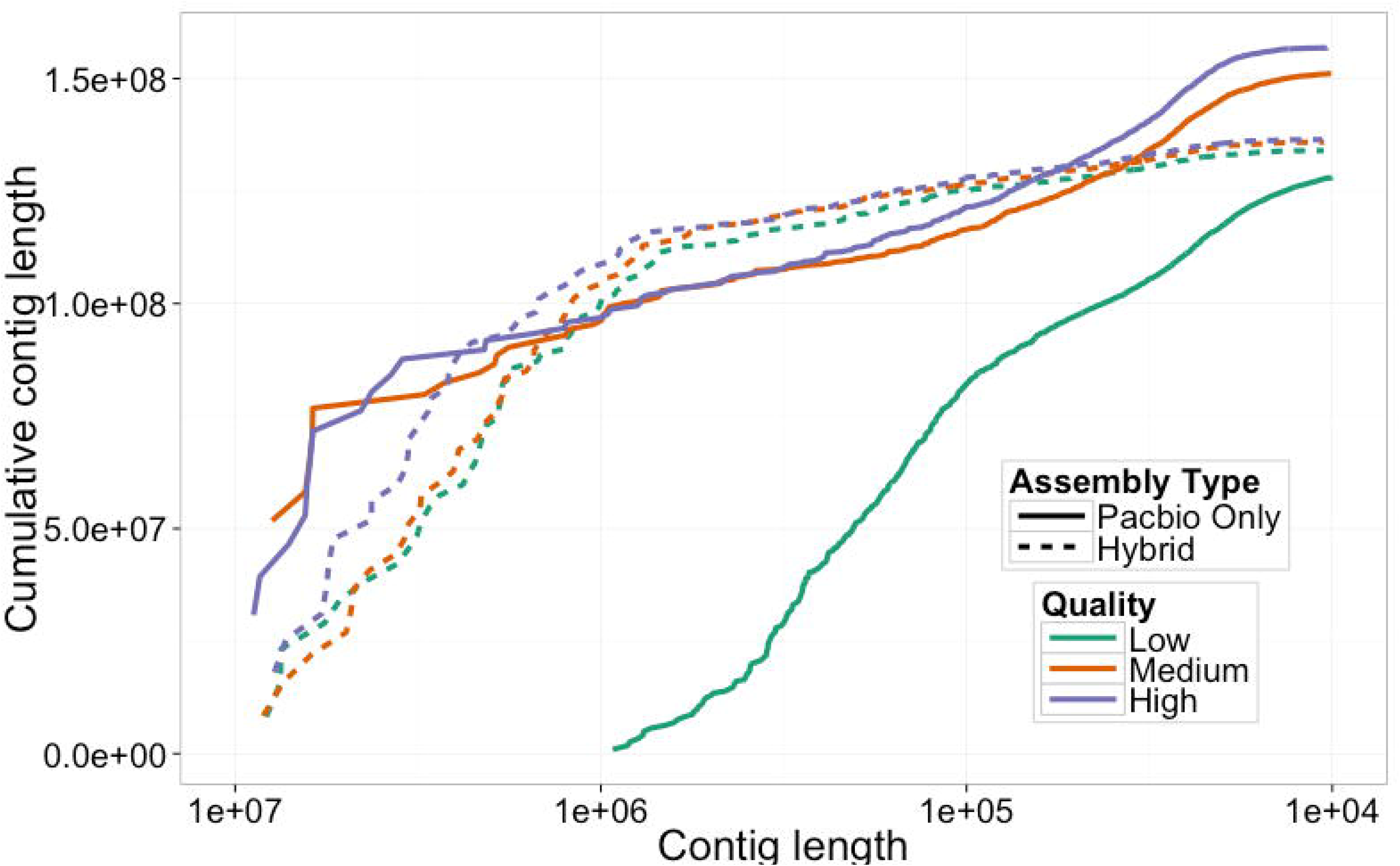
As in Figure 2a, a plot of cumulative length distribution. These curves represent the cumulative length distribution of final assemblies using low, medium, and high quality selected reads using either PacBio only assembly or hybrid assembly.

### Merging of human assemblies of the CHM1 cell line

One challenge in a study of this type is determining whether mergering performed on a very different genome, like that *of Homo sapiens,* would performed as well as on *D. melanogaster.* To do this, we used publicly available sequence data and assemblies for the human hydatidiform mole (CHM1 (36)) to generate a merged assembly for *H. sapiens,* both to gauge the performance of *quickmerge* on a different species than it was developed on, and to observe its performance on a larger and more repetitive genome (the human genome is ~3.2Gb, approximately 25X the size of the *D. melanogaster* genome).

Of the available CHM1 data, we chose to re-use the data used in Berlin et al. 2015 (14) (the P5C3 chemistry). We ran our genome assembly pipeline on the 30X longest reads of PacBio data from the 54X in the CHM1 dataset, plus 40.66x of publicly available human CHM1 Illumina data (NCBI accession: PRJNA176729). The hybrid assembly produced an NG50 of 2.4Mb, which is in line with the results observed in Fig. 5. Along with this, we used the PacBio assembly contigs produced by Berlin et al. 2015, which had an NG50 of 4.1Mb. We merged the two assemblies with more strict parameters because of the larger genome size: we set HCO to 15, c to 5, and l to 5Mb. Merging the two assemblies produced a final assembly NG50 of 8.85Mb, a substantial improvement upon the PacBio only assembly. This more than doubling of NG50 is in line with our expectations based on the *D. melanogaster* results; all available data indicate that this pipeline improves contiguity for CHM1 to the same extent that it does for the *D. melanogaster* ISO1 strain. We did not polish this assembly with Quiver and Pilon due to computational constraints, but it stands to reason that the gains vis-à-vis SNP and indel rates would be similar between human and *D. melanogaster.* In order to evaluate misassemblies, we produced a MUMmer dnadiff report by comparing the PacBio only, DBG2OLC, and merged assemblies to the most recent and highest contiguity CHM1 PacBio only assembly available (GenBank accession number: GCA_001420765.1). The results show that the large increase in contiguity is not a consequence of merging induced misassembly, mirroring the results in *Drosophila* (Supplementary Fig. 12). Additionally, we generated MUMmer dot plots that indicated that contig orientation and ordering were correct, with the exception of some inversions and translocations that were inherited from the component assemblies (Supplementary Fig. 7). While we attempted to run the Quast and GAGE assessment pipelines on the human assemblies, we found that, in all cases, the programs either crashed or failed to finish successfully in a reasonable time frame.

## Discussion

Genome assembly projects must balance cost against genome contiguity and quality (4). Self correction and assembly using only long reads clearly produces complete and contiguous genomes (Fig. 5; supplementary Table 1). However, it is often impractical to collect the quantity of PacBio sequence data (>50X) necessary for high quality self correction either because of price or because of scarcity of appropriate biological material, especially when assembling very large genomes. For example, at least 40 μg of high quality genomic DNA is required for us to generate 1.5 μg of PacBio library when we use two rounds of size selection in the library preparation protocol. A 1.5 μg library produces, on average, 15-20 Gb of long DNA molecules. This dramatic loss of DNA during library preparation limits the amount of PacBio data that can be obtained for a given quantity of source tissue. When a project is limited by cost or tissue availability, a hybrid approach using a mix of short and long read sequences is an alternative to self corrected long read sequences.

Our results show that when 67.4X of 100bp paired end Illumina reads is used in combination with 10X -30X of PacBio sequences, reasonably high quality hybrid assemblies can be obtained, with approximately 30X of PacBio sequences yielding the best assembly. In fact, as our results show, a 30X hybrid assembly is less fragmented and higher quality than even a 50X self-corrected assembly (Fig. 5). However, our results also show that with the same long molecule data, PacBio only and hybrid assemblies often assemble complementary regions of the genome. The implication here, that different assemblers are joining complementary contigs, suggesting that future assemblers could generate higher quality assemblies with modest coverage data. The merging of a PacBio only and a hybrid assembly results in a better assembly than either of the two alone (Figure 5, supplementary table 1), regardless of the total amount of long molecule sequences (≥30X) used. Thus, projects for which ≥30X of single molecule sequence can be generated are well-served by collecting an additional 50-100X of Illumina data. These data can then be used to generate both a self-corrected assembly and a hybrid assembly, which can then be merged to obtain an assembly of comparable contiguity to PacBio only assemblies using twice the amount of PacBio data (Fig. 5). This merged assembly approach produced the highest NG50 of any assembly at all coverage levels at which it could be tested, with little or no tradeoff in base accuracy or misassemblies (Supplementary Fig. 8-10).

Nonetheless, it is clear that the tools available for genomic assembly have inherent technical limitations: DBG2OLC assembly contiguity asymptotes as PacBio read coverage passes about 30X, and the PBcR pipeline produces the best assembly when the longest reads that make up 40X (of genome size) of data are corrected and only the longest 25X from the corrected sequences are assembled (14). Indeed, when coverage greater than 25X is used for PacBio only assembly, there is a real loss of assembly quality as coverage increases (data not shown). This may be because an increase in coverage leads to the stochastic accumulation of contradictory reads that cannot be easily reconciled, a limitation of the overlap-layout-consensus (OLC) algorithm used in assembling the long reads(2,37).

Long read sequencing technologies, such as those offered by PacBio, Oxford Nanopore (38), and Illumina TrueSeq (39) promise to improve the quality of *de novo* genome assemblies substantially. However, as we have shown using PacBio sequences as an example, not all long read data is equally useful when assembling genomes. We provide empirical validation, perhaps for the first time, of length and quality on assembly contiguity. Additionally, our results provide a novel insight: high throughput short reads can still be useful in improving contiguity of assemblies created with long reads, even when long read coverage is high. In light of our results, we have a compiled a list of best practices for DNA isolation, sequencing, and assembly (Supplementary Fig. 15 and Supplementary Fig. 16). Particularly important for DNA isolation is quality control of read length via pulsed field gel electrophoresis. Regarding assembly, we recommend that researchers obtain between 50x and 100x Illumina sequence. Next, researchers must determine how much long molecule coverage to obtain: between 25x and 35x, or greater than 35x. With coverage below 35X, PacBio only methods often fail to assemble, and produce low contiguity when they do assemble, and thus, we can only confidently recommend a hybrid assembly. Above 35X, we recommend meta assembly of a hybrid and a PacBio only assembly. In this case, we recommend downsampling to the 30X longest PacBio reads when generating the hybrid assembly because hybrid assembly contiguity decreases above this coverage level, but this has not been extensively tested. We show that this approach is effective both in *Drosophila* and human genomes, which differ in size and extent of repetitive regions.

One challenge in assembly is posed by analyzing data from heterozygous individuals. Heterozygosity is known to make assembly more challenging (5). All of the data evaluated in this study were produced from either isogenic or highly inbred populations (*Drosophila)* or from a single haploid cell line (human CHM1). Because there is not a comparable dataset available that was produced using heterozygous individuals, we cannot test the effect of heterozygosity on assembly quality. That said, some assemblers (*Platanus* (24) and *Falcon* (https://github.com/PacificBiosciences/FALCON) were designed to produce diploid assemblies from heterozygous sequence data (5). It stands to reason that substituting *Falcon* in the place of *PBcR* in this pipeline could improve assembly quality for highly heterozygous samples, but that claim will require further testing.

The recent rapid development of short read sequencing technology has fostered an explosion of genome sequencing. However, as a result of the cost effectiveness and concomitant popularity of short read technologies, the average quality and contiguity of published genomes has plummeted (40). Indeed, short read sequences are poorly suited to the task of assembly, especially when compared with long molecule alternatives. While long molecule sequencing has rekindled the promise of high quality reference genomes for any organism, it is currently substantially more expensive than short read alternatives. In order to mitigate uncertainties inherent in adopting this technology, we have outlined the most salient features to consider when planning a genome assembly project. We have recommended effective DNA isolation and preparation practices that result in long reads that take advantage of what the PacBio technology has to offer. We have also provided a guide for assembly that leads to extremely contiguous genomes even when circumstances prevent the collection of large quantities of long molecule sequence data recommended by current methods.

## Acknowledgement

The authors would like to thank Stephen Richards for sharing the length distribution from *Drosophila pseudoobscura* Pacific Biosciences data and Sergey Koren and Brian Walenz for their assistance with wgs. We would also like to thank Melanie Oakes and Valentina Ciobanu for assistance with sequencing.

